# A Deep learning approach for Pan-Renal Cell Carcinoma classification and survival prediction from histopathology images

**DOI:** 10.1101/559401

**Authors:** Sairam Tabibu, P.K. Vinod, C.V. Jawahar

**Affiliations:** Center for Visual Information Technology, IIIT, Hyderabad, India; Center for Computational Natural Sciences and Bioinformatics, IIIT Hyderabad, Hyderabad, India

## Abstract

Histopathological images contain morphological markers of disease progression that have diagnostic and predictive values. However, complex morphological information remains unutilized in unaided approach to histopathology. In this study, we demonstrate how deep learning framework can be used for an automatic classification of Renal Cell Carcinoma (RCC) subtypes, and for identification of features that predict survival outcome from digital histopathological images. Convolutional neural networks (CNN’s) trained on whole-slide images distinguish clear cell and chromophobe RCC from normal tissue with a classification accuracy of 93.39 % and 87.34 %, respectively. Further, a CNN trained to distinguish clear cell, chromophobe and papillary RCC achieves a classification accuracy of 92.61 %. Here, we introduced a novel support vector machine based method to deal with data imbalance in multi-class classification to improve the accuracy. Finally, we extracted the morphological features from high probability tumor regions identified by the CNN to predict patient survival outcome of most common clear cell RCC. The generated risk index based on both tumor shape and nuclei features are significantly associated with patient survival outcome. These results highlight that deep learning can play a role in both cancer diagnosis and prognosis.

## Introduction

Kidney Cancer accounts for nearly 3.8% of adult cancers and is among the 10 most common cancers in both men and women. Recent estimate by American Cancer Society indicates that 63340 new cases and 14970 death will occur in 2018^1^. Renal Cell Carcinoma (RCC) is the most common (85%) malignant tumor of kidney and is a heterogeneous group of tumors with different histology, molecular characteristics, clinical outcomes and responses to therapy^2^. The major subtypes of RCC are clear cell, papillary and chromophobe accounting for 70-80%, 14-17% and 4-8% of RCC, respectively^3, 4^.

Clear cell RCC (KIRC) and papillary RCC (KIRP) originate from cells in the proximal convoluted tubules of the nephron^5^. KIRC is characterized with loss of chromosome 3p and mutation of the von Hippel–Lindau (VHL) gene while KIRP is characterized by trisomy of chromosomes and loss of chromosome 9p^6, 7^. Chromophobe RCC (KICH) originates from intercalated cells in the distal convoluted tubules and is characterized by loss of chromosomes^8^. KIRC patients have an overall 5 year survival rate of 55-60%^9–11^ whereas for KIRP patients, it vary from 80-90%^12, 13^ and for KICH patients, it is 90%^14^. Due to these distinct biological and clinical behavior of subtypes, accurate detection of RCC and its subtypes is vital for the clinical management of patients.

RCC subtypes can be detected radiologically based on degree of tumor enhancement on multidetector computed tomography (CT) or magnetic resonance imaging (MRI)^15–19^. Further, the microscopic examination of Hematoxylin and Eosin (H & E) stained slides of biopsies continues to be a valuable tool for pathologist and clinicians^20^. Histological images contain markers of disease progression and phenotypic information that can have diagnostic and predictive values. Major limitations in the examination of H & E images by pathologists are inter-observer discordance and time required to diagnose. The pathologist follows a simple decision tree approaches utilizing only a fraction of available information. Complex morphological information remains unutilized in unaided approach to histopathology.

The Cancer Genome Atlas (TCGA)^21, 22^ project has resulted in the creation of large repositories of digital H & E whole-slide images (WSI) of RCC. These images are acquired at 20x and 40x magnifications with size varying from 10000-100000 pixels, which are visually tricky to analyze and interpret accurately. Further, the high-resolution WSI also poses a challenge to develop computational model for classification. An automated system to detect RCC and its subtypes is still not available. Cheng *et.al.* (2017)^23^ extracted image features from TCGA tumor slides to develop a lasso-regularized Cox model for predicting survival of KIRC patients.

In recent years, deep learning techniques especially convolutional neural networks (CNN’s) have significantly improved accuracy on a wide range of computer vision tasks such as image recognition^24–26^, object detection^27, 28^ and semantic segmentation^29, 30^. Further, CNN’s have also been successful in capturing the complex tissue patterns and have been widely used in biomedical imaging for segmentation as well as for classification tasks in cancers such as breast^31–33^, lung^34–37^ and prostate^38^.

In this study, we demonstrate how deep learning can be applied to histopathological images of RCC. An automatic system for pan-RCC classification and survival prediction tasks was developed from WSI of TCGA. CNN’s trained on WSI classify each RCC subtype from normal tissues and three RCC subtypes, and extract features for survival prediction. In that process, we also characterized the high probability cancerous regions and tissue of origin differences, and came up with a strategy for overcoming data imbalance in multi-class classification problems using histopathological images.

## Results

### CNN model to distinguish RCC from normal tissue using histopathology images

TCGA slide images of RCC and normal tissues were split into 70-15-15 % for training, validation and testing (**Supplementary Table 1-3**). Image patches of size 512 × 512 were extracted from the slides of both 20x and 40x resolution with 50% overlap (**Figure 1**). The background patches were removed using pixel thresholding (See Methods Section). We used the pre-trained Resnet 18 and Resnet 34 by replacing the last layers with just 2 output layers and fine-tuned it on RCC data. We terminated the training after 30 epochs when the validation accuracy failed to improve. The images were processed patch-wise as well as slide-wise. We trained the model on only KIRC and KICH due to lack of sufficient normal tissue samples for KIRP. We obtained 93.39 % and 87.34 % **(Table 1 and 2)** patch-wise accuracy on the test set for KIRC and KICH, respectively. To assess this accuracy, we performed a slide-wise analysis by counting the percentage of positively classified patches in a slide. We achieved a best Area Under Curve (AUC) of 0.98 for KIRC and 0.95 for KICH using Resnet 34 **(Table 2)**.

**Figure 1.**
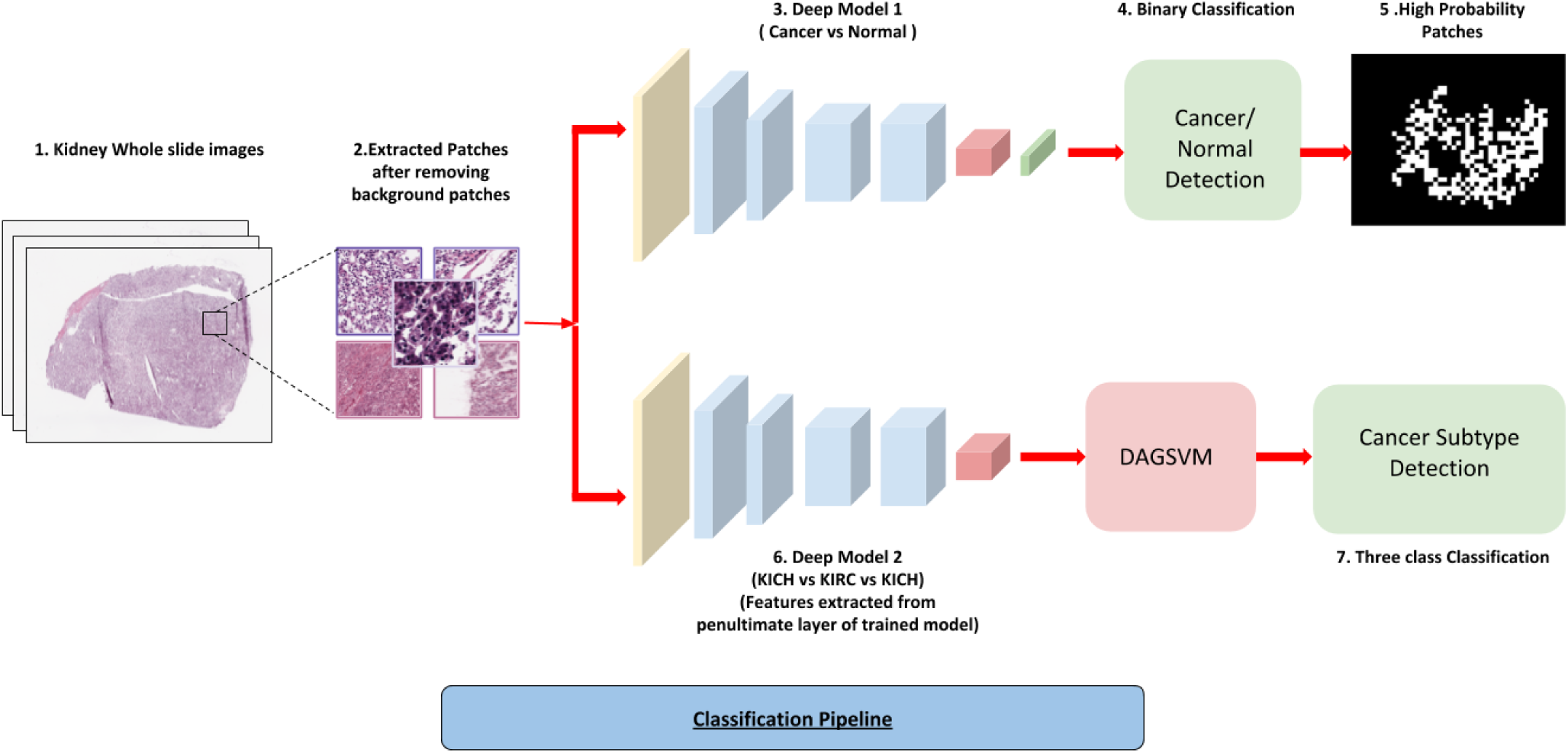
Cancer Classification Pipeline. **1.** Kidney Whole slide images. **2.** 512*512 patches extracted from images with 50% overlap and background removed using pixel thresholding. **3.** Patches from normal and cancerous slides fed to the deep network. **4.** Patches classified as cancerous or non-cancerous. **5.** High probability patches identified by the trained network and binary mask is applied. **6.** The patches from three subtypes used to train a similar deep architecture for a three-way classification. **7.** Features extracted from the penultimate layer of the network and fed to DAGSVM and a three way classification is performed by it.

**Table 1.**
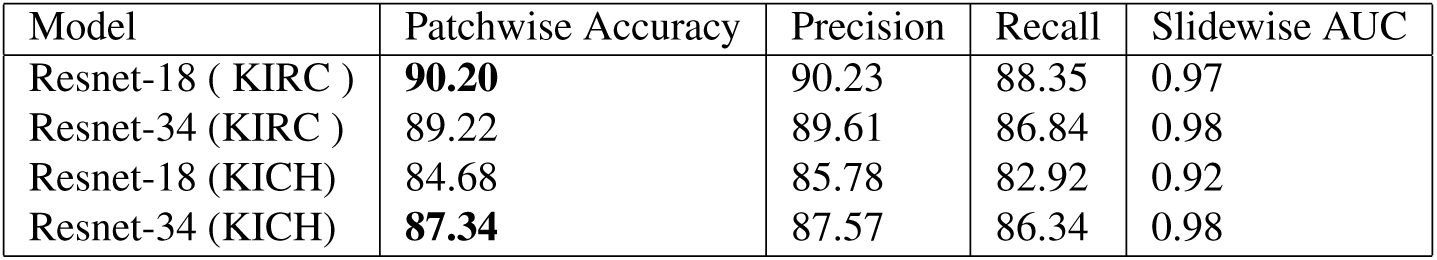
Cancer/Normal Classification (20x resolution)

| Model | Patchwise Accuracy | Precision | Recall | Slidewise AUC |
| --- | --- | --- | --- | --- |
| Resnet-18 ( KIRC ) | **90.20** | 90.23 | 88.35 | 0.97 |
| Resnet-34 (KIRC ) | 89.22 | 89.61 | 86.84 | 0.98 |
| Resnet-18 (KICH) | 84.68 | 85.78 | 82.92 | 0.92 |
| Resnet-34 (KICH) | **87.34** | 87.57 | 86.34 | 0.98 |

**Table 2.** Cancer/Normal Classification (40x resolution)

| Model | Patchwise Accuracy | Precision | Recall | Slidewise AUC |
| --- | --- | --- | --- | --- |
| Resnet-18 ( KIRC ) | **93.39** | 93.41 | 92.95 | 0.99 |
| Resnet-34 (KIRC ) | 93.62 | 93.47 | 93.37 | 0.99 |
| Resnet-18 (KICH) | 79.09 | 79.61 | 80.06 | 0.95 |
| Resnet-34 (KICH) | **80.57** | 80.65 | 81.17 | 0.95 |

### RCC subtype detection using CNN

A challenging task is to distinguish the subtypes of RCC: KIRC, KIRP and KICH. We used the deep learning framework to distinguish these three subtypes (**Figure 1**). However, there exists a major class imbalance with 43 % KIRC, 14 % KIRP and 43% KICH images. First we tested whether the CNN can distinguish these subtypes irrespective of the data imbalance. We extracted the 512 × 512 patches from 20x resolution for each subtype and performed a three class classification using Resnet-18 and Resnet-34. We obtained a patchwise accuracy of 87.69 % and a micro-average AUC of 0.91 **(Table 3)**. However, the recall score is in the range of 83 % suggesting that certain classes are misclassified. Individual analysis of the classification scores on each subtypes **(Supplementary table 4)** revealed that the CNN performs better on subtypes KIRC and KICH but not on KIRP (70%) which can be attributed to the severe data imbalance within the subtypes.

**Table 3.** Cancer Subtype Classification

| Model | Patchwise Accuracy | Precision | Recall | Slidewise AUC |
| --- | --- | --- | --- | --- |
| Resnet-18 | 87.69 | 88.82 | 83.66 | 0.88 |
| Resnet-34 | 86.19 | 88.30 | 83.18 | 0.88 |
| Resnet-18 + DAGSVM | **92.62** | 90.78 | 89.07 | **0.93** |
| Resnet-34 + DAGSVM | **91.96** | 88.94 | 87.92 | **0.93** |

We used a DAG-SVM method^39^ to deal with this data imbalance method without making any changes to the deep learning architecture. In this framework, the N-class classification problem is divided into N(N-1)/2 binary classification problem by replacing the softmax layer of deep architecture by binary classifiers for each pair of classes arranged in a directed acyclic graph (DAG) structure (**Figure 3**). Such division enables the network to focus on individual binary problems which are relatively easier to tackle as well as to learn more pairwise discriminative features (See Methods Section). Using the representation matrix obtained by the CNN from the penultimate layer of the CNN for patch images, we trained the DAG-SVM and performed the subtype classification. We obtained a substantial increase in the patch-wise accuracy by 5% **(Table 3)**. The increase in the micro-average AUC is 0.03 whereas macro-average AUC is 0.05. This demonstrates that the combined CNN and DAG-SVM model substantially improves the performance of discriminating the subtypes even in the given limitation of data imbalance. This is also evident from the increase in the classification score of the subtype KIRP (10%) **(Supplementary Table 5)**.

### Prediction of Tumor Areas and the Survival outcome for KIRC

Using the CNN model that distinguishes RCC from normal tissue, high probability tumor patches were identified and a probability heat map was generated for the WSI. Based on the tumor region, several tumor and nuclei shape features such as area, perimeter etc. were extracted from the cancer slide images of each patient **(Figure 2)**. The risk index of each patients was calculated using lasso-regularized COX model for each feature, and validated using two-level cross validation strategy (see methods).

**Figure 2.**
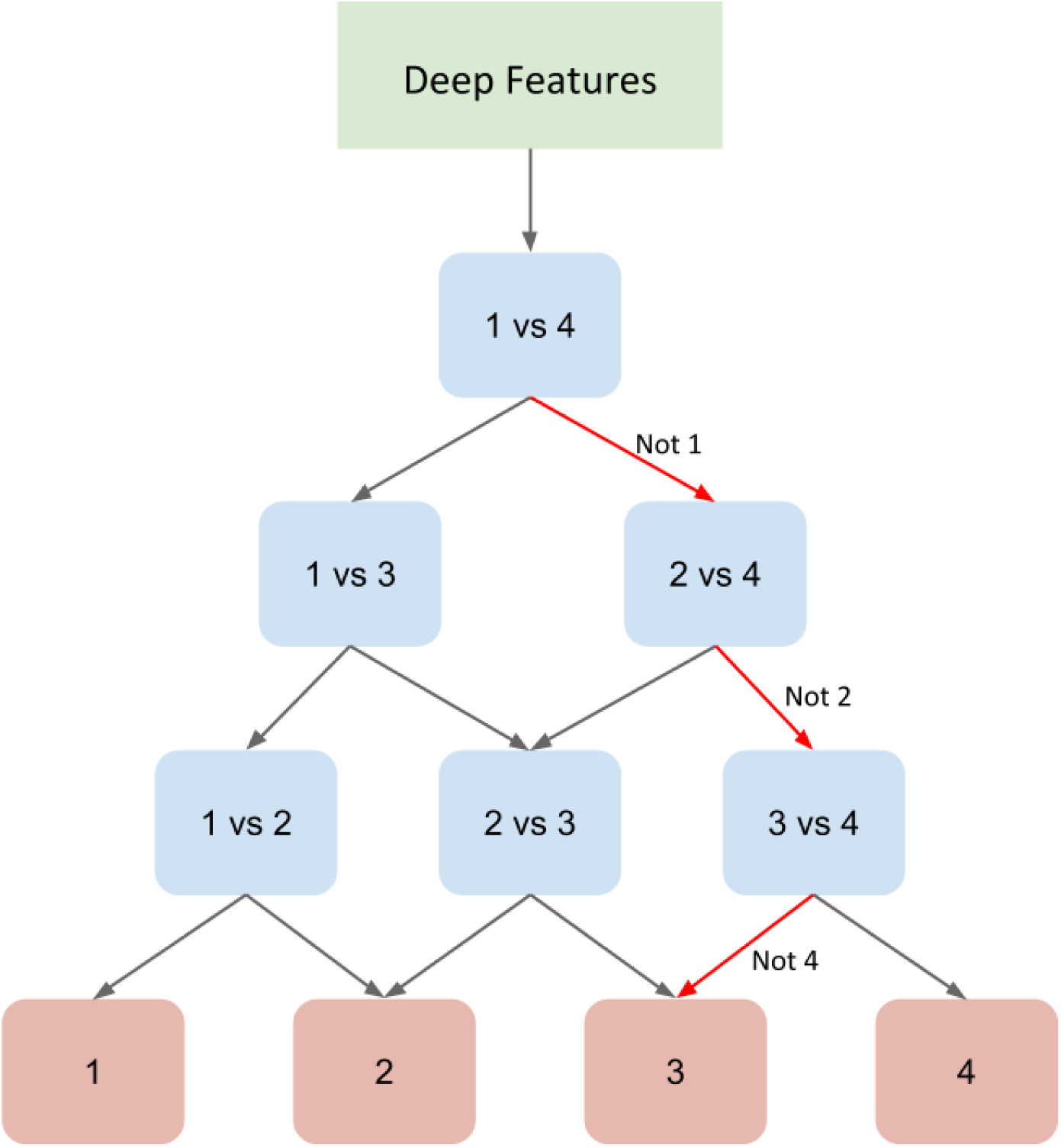
Directed Acyclic Graph SVM (DAGSVM) Architecture of the model for a four-class problem where features learned by the deep network are used to train all the classifiers. Each node is a binary classifier for a pair of classes.

**Figure 3.**
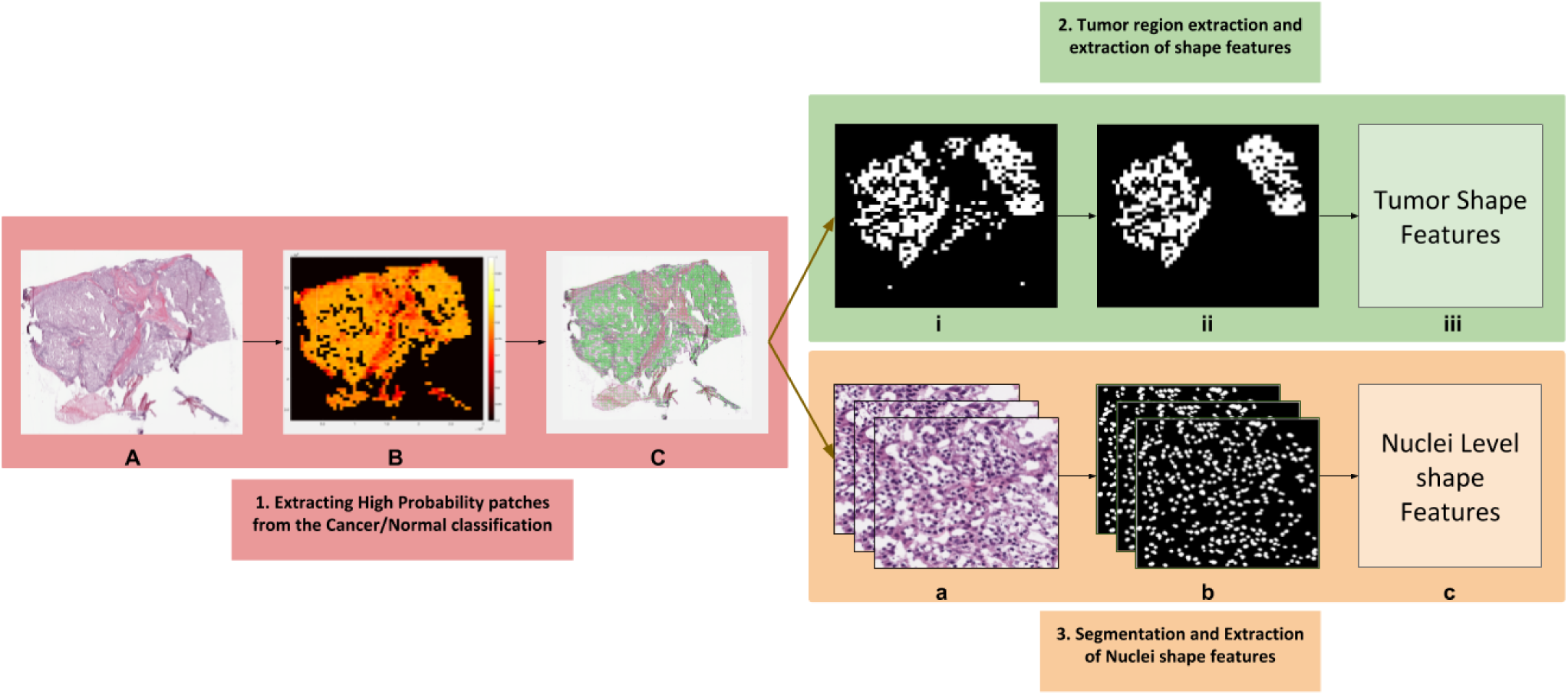
Shape Features Extraction Pipeline. **A.** Tissue slide image. **B.** Heatmap generated after each patch is fed through the deep network. **C.** High probability patches are highlighted. **i.** Binary mask generated for high probability patches. **ii.**Small patches removed using morphological operations. **iii.** Tumor shape features extracted from final image. **a.** High probability patches accumulated. **b.** Nuclei segmented from each patch. **c.** Nuclei shape features extracted from each patch.

**Figure 4** shows the survival curves of low and high risk groups for different features. We found that 13 tumor shape features and 6 nuclei shape features are significantly (p value less than 0.05) associated with the patient survival **(Table 4)**. Tumor shape features include total area (p value = 1.5e-6, HR = 2.398) and total perimeter (p value = 1.49e-5, HR = 2.485), and nuclei shape features include total convex area (p value = 2.2e-7, HR = 2.576) and total major axis (p = 0.000614, HR = 2.252) **(Figure 4 A, 4 B and 4 C)**. Most predictive individual features were selected for integrative analysis to stratify patients into low risk and high risk groups using Lasso-Cox model. We found significant (p = 3.68e-6) association between combined image features and survival outcome **(Figure 4 D)**. Multivariate analysis was performed with predicted risk indices, age, gender, grade and stage as covariates **(Table 5)**. We observed that the survival outcome significantly depends on the predicted risk indices and stages of tumor. This suggest that tumor and cell nuclei shape features identified in this study can be used as a prognostic factor to predict the survival outcome of KIRC patients.

**Figure 4.**
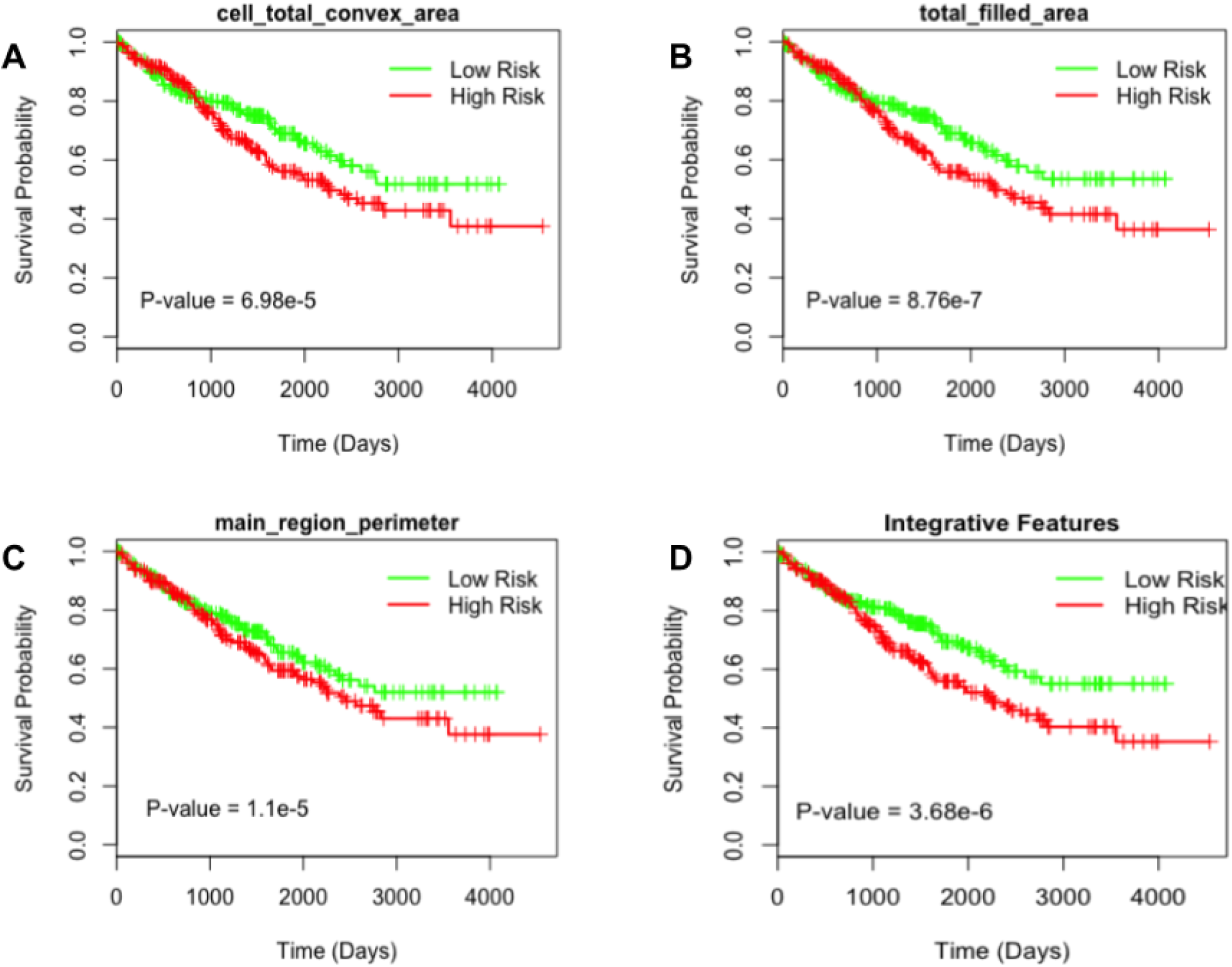
Tumor shape features and cell shape features predict survival outcome of ccRCC patients.

**Table 4.** Tumor shape and Nuclei Features and their log Rank test scores

| Feature type | Image Feature | P-value |
| --- | --- | --- |
| Tumor Shape Features | Total Area | 1.5e-6 |
|  | Total convex area | 2.2e-7 |
|  | Total perimeter | 1.49e-5 |
|  | Total filled area | 8.76e-7 |
|  | Total major Axis | 0.000614 |
|  | Total minor axis | 0.000572 |
|  | Total perimeter by Area | 0.00139 |
|  | Main region area | 0.0042 |
|  | Main region convex area | 1.58e-5 |
|  | Main region perimeter | 1.1e-5 |
|  | Main region perimeter by area | 4.29e-5 |
|  | Main region major axis | 6.09e-5 |
|  | Main region minor axis | 0.00161 |
| Nuclei shape features | total area | 0.000106 |
|  | Total convex area | 6.98e-5 |
|  | Total perimeter | 0.00327 |
|  | Total filled area | 0.000115 |
|  | Total major axis | 0.00251 |
|  | Total minor axis | 0.00748 |
| Integrative model |  | 3.68e-6 |

**Table 5.** Multivariate analysis of predicted risk indices in TCGA.

| Variable | HR(95% CI) | p-value |
| --- | --- | --- |
| Lasso Cox | 2.265(1.5343-3.343) | 3.87e-5 |
| Age | 1(0.999-1.0001) | 3.71e-6 |
| Gender | 1.095(0.7870-1.524) | 0.590 |
| Stage | 1.733(1.4968-2.007) | 2.01e-13 |
| Grade | 1.106(0.9369-1.306) | 0.233 |

## Discussion

In this study, we developed an automatic system that can distinguish RCC from normal tissue and RCC subtypes using histopathology images. To the best of our knowledge, this is the first study to perform classification of RCC subtypes using deep learning framework. CNN’s successfully identified the cancerous tissue patterns as well as the inherent texture differences among RCC subtypes. We also introduced a method to deal with data imbalance which can be generalized to other cancer classification tasks. Finally, we developed a prognostic model based on the regions detected by the CNN to predict survival outcome. We used a two-level cross validation technique to validate our model.

Due to the enormous amount of information contained in the WSI, the histopathological analysis is a laborious and time intensive task for the pathologist. With the growing number of cancer cases, fast and precise assessment is not feasible. Our study addresses this limitation by providing computational interpretation of WSI with high AUCs in the testing phase of various classification tasks. Since the pipeline was developed on the WSI of cancer, it eliminates the requirement of annotating tumor regions for the training purpose and provides a strategy that can used to study WSI of other tumors. Further, a high patch-wise AUC obtained for distinguishing tumor from normal tissue suggests that tumor regions are not localized and are spread across WSI. Our analysis also showed that a better performance is obtained for KIRC when trained on 40x images and for KICH when trained on 20x images **(Table 1 and 2)**. These results suggests the existence of complex morphological distinguishing features between cancers.

Subtypes of RCC typically have varying morphological patterns and nuclear/cytoplasmic features at the microscopic level^40^. Different cases of KIRC have mixture of cells with clear cytoplasm and granular eosinophilic cytoplasm, pseudopapillary pattern, rich vascularity and areas of hemorrhage. KICH has polygonal cells with prominent cell membranes, cells of different size with either eosinophilic or foamy cytoplasm and incomplete vascular septae. KIRC has predominantly papillary growth pattern, foamy macrophages and cuboidal cells. However, there are instances were histopathology images are not conclusive leading to re-classification of some KIRC samples as KICH during the histological review^41^.

The CNN model is able to distinguish these subtypes by capturing their unique characteristics with high AUC of 0.93 **(Table 3)**. Further, we also tested whether the inherent difference in the texture of tissue of origin contribute to classification of subtypes by using normal tissue samples as a test set to RCC subtype model. We obtained subtypes classification with AUC of 0.85 **(Supplementary Table 6)**. Together, these analyses suggest that both tumor regions and the inherent texture difference contributes to subtype classification. Texture classification methods^42^ can be further used to visualize the difference **(Figure 5)** and study their characteristics. Further, for subtype detection, we introduced a DAG-SVM classifier on the top of the deep network without introducing any change to the structure of the Network. This helped to break the multiclass classification task into multiple binary classification tasks which not only improved the performance of the model but also helped to deal with data imbalance.

**Figure 5.**
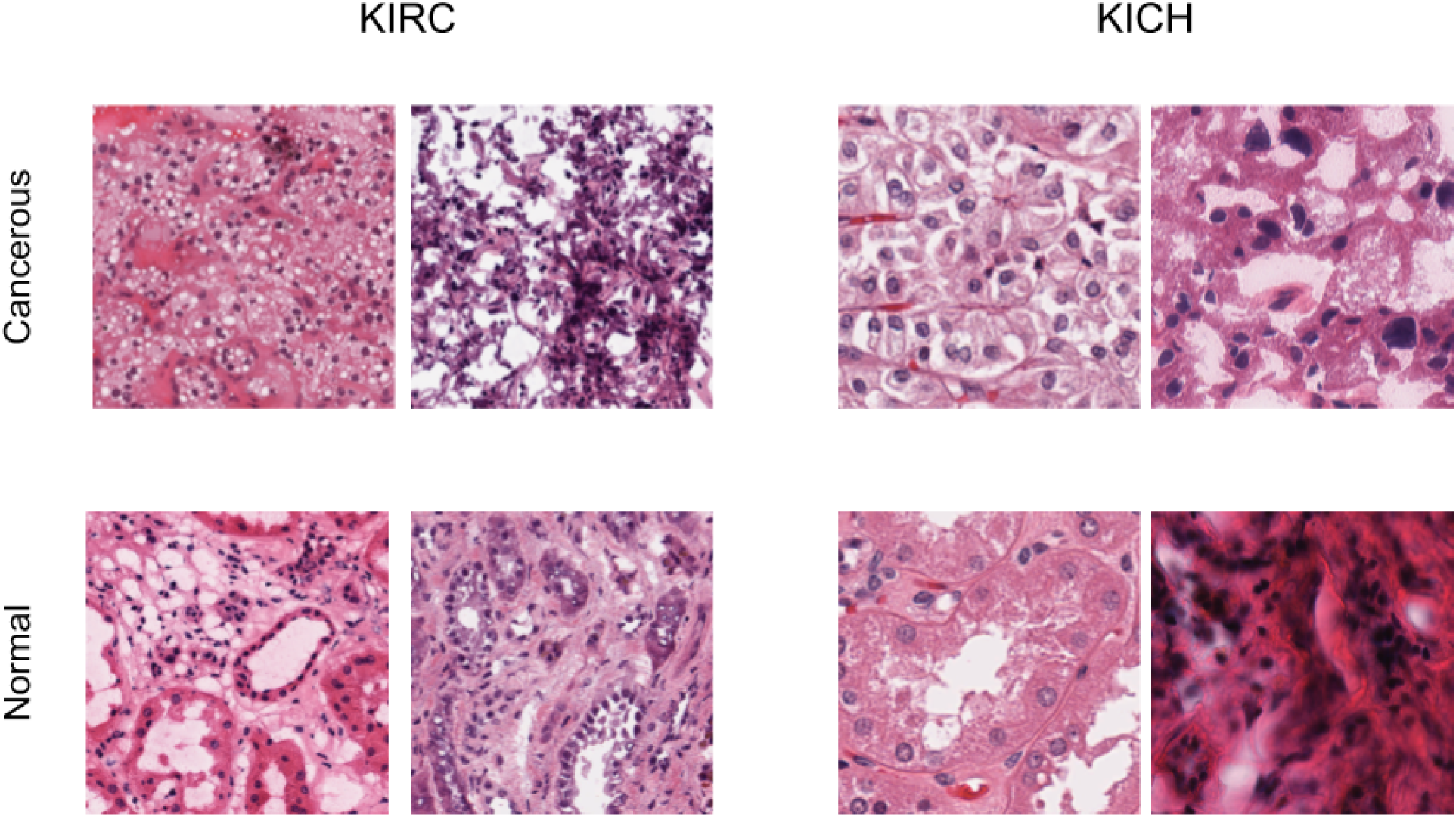
Visualization of Cancerous and Normal tissue regions at 40x resolution.

Finally, we also demonstrated the reliability of features extracted from tumor slide images in predicting the survival outcome. We showed that both tumor shape and nuclei features from the high probability regions identified by CNN are significantly associated with survival outcome of KIRC. Image features such as ratio of perimeter squared by area, which is a quantification of tumor surface^35^, is correlated with survival outcome. A major finding that tumor shape features also predict survival extends the previous studies that uses genomic and nuclei features to predict survival of KIRC. It is also consistent with observation that tumor shape features can be used to predict the survival outcome of lung patients^35^. However, this model is developed using annotated images by pathologist, which in our case is done independent of any human bias.

The lack of data and the class imbalance are major challenges in building survival models of other subtypes. Firstly, there is dearth of KIRP-matched normal slide images that can be used to develop a model to distinguish tumor from normal tissue and to identify high probability cancerous regions. Secondly, since the ratio of events (death) to total number of patients is few for KICH and KIRP (less than 15%) compared to KIRC (around 40%), the survival prediction for KICH and KIRP is affected. Further, subtypes such as renal oncocytoma and collecting duct (duct Bellini) of RCC have the rarest chance to occur (less than 1%) and therefore data is scarce to develop model including rare subtypes. On the other hand, we made several attempts to use only tumor slide images of KIRP to classify different tumor stages and grades, which did not yield satisfactory performance. Further study in this direction is required to see whether improvements in deep learning framework can help to detect the subtle differences between early and late stages of RCC and to integrate genomic features to improve the performance. Studies to understand the complex relationship between tissue histology and genetics can provide useful insights for the the diagnosis and treatment.

## Methods

### Dataset and Image Processing

The whole slide images and clinical information were downloaded from TCGA data portal (https://gdc.cancer.gov/). Slides with reading and compatibility issues were removed (972 slides in total were removed from the whole dataset). We selected 1027 (KIRC), 303 (KIRP), and 254 (KICH) tumor slide images for our study. Further, corresponding 379, 47 and 83 normal tissue slide images for each subtype were selected. 512*512 sized tiles were extracted with 50% overlap ensuring multiple viewpoints within the tissue, at a magnification of 20x and 40x. For subtype classification, patches from 20x magnification were used. The patches were removed if the mean intensity of 50% pixel values was larger than a threshold (in our case 210 for RGB channels).

### Training Architecture and Framework

The Resnet 18 and 34 architectures^24^ pre-trained on Imagenet dataset were used with input resized from 512*512 to 224*224. The final linear layer was modified to 2 for cancer/normal detection and 3 for subtype detection. The network was fine-tuned with stochastic gradient optimizer and cross entropy was used as the loss function. To introduce generalization, we also used data augmentation techniques (random horizontal flip and random Crop) while training. The images before being fed to the network were normalized using mean and standard deviation which was calculated on the training set. Batch size was set to 128 with learning rate 1e-5 and decay according to the validation error. The average number of epochs for both experiments lies in between 3-40 epochs. The training was terminated when the validation accuracy did not vary much for 4-5 epochs. Tests were implemented using Pytorch library^43^. For the 3-class classification problem in the subtype detection, we used a DAG-SVM on the top of the fully connected layer of the CNN to improve the multiclass accuracy. We removed the final softmax layer from the trained model and use the learned representation given by the CNN (which is essentially a 512*1 vector) to train the DAG-SVM. Linear kernel was used while training the SVMs.

### Mapping high probability regions and survival analysis

The survival analysis was performed on KIRC WSI’s **(Supplementary Table 7)** The probability score for each patch was calculated by passing them through the KIRC CNN model (normal/cancer) trained on patches in 40x resolution, and a heatmap was constructed for each slide**(Supplementary Figure 1&2)**. Taking a suitable threshold, the high probability regions were highlighted using binary masking. To remove the effects of very small patches, tissue regions which were less than 1/3rd of the main region were removed. 15 shape features including region area, convex area, perimeter, solidity, and eccentricity were extracted from the high probability patches. Further, the nuclei were segmented^44^ and 7 nuclei shape features were also extracted **(Supplementary Table 8)**. If a patient had one or more slides the values were averaged and patient level features were aggregated. We used these morphological features to build a Lasso regularized COX model. We estimated the risk index of each patient based on each morphological feature. A two level cross validation strategy was used to validate our model^23^. Each patient was kept aside as a test case and the rest of the samples were used to train the COX model using a 10 fold cross validation. The model was then used to stratify the left out patient into either high risk or low risk group using the median score obtained from the training set. This process was repeated until each patient was used as a test set. From the finally obtained groups and survival indices, p-values were calculated using log-rank test. We used Kaplan-Meier plots to visualize the survival curves of low and high risk groups. The survival analysis was performed using R packages “survival” and “glmnet”^45, 46^.

## Author contributions statement

S.T. developed the models for various tasks. S.T., P.K.V. and C.V.J analyzed and discussed the data. S.T., P.K.V. and C.V.J wrote the manuscript. P.K.V. and C.V.J. conceived the study and were in charge of overall direction and planning.

## Additional information

The authors declare that they have no competing financial interests

