## Supplementary information for "A Deep learning approach for Pan-Renal Cell Carcinoma classification and survival prediction from histopathology images"

Supplementary Table 1: Number of slides and images for RCC normal and Cancerous tissue for 20X and 40X respectively for KIRC.

| Resolution | Tissue | Training | Validation | Testing |
|---|---|---|---|---|
| 20X | Cancerous | 719(213302) | 154(55463) | 154(42907) |
| 20X | Normal | 379(115856) | 81(23434) | 81(23401) |
| 40X | Cancerous | 719(672586) | 154(163093) | 154(130399) |
| 40X | Normal | 379(428719) | 81(88406) | 81(92581) |

Supplementary Table 2: Number of slides and images for RCC normal and Cancerous tissue for 20X and 40X respectively for KIRP.

| Resolution | Tissue | Training | Validation | Testing |
|---|---|---|---|---|
| 20X | Cancerous | 179(74626) | 39(23613) | 38(13577) |
| 20X | Normal | 33(6102) | 7(1285) | 7(908) |
| 40X | Cancerous | 179(235858) | 39(83855) | 38(81267) |
| 40X | Normal | 33(21614) | 7(4365) | 7(5418) |

Supplementary Table 3: Number of slides and images for RCC normal and Cancerous tissue for 20X and 40X respectively for KICH.

| Resolution | Tissue | Training | Validation | Testing |
|---|---|---|---|---|
| 20X | Cancerous | 106(202912) | 23(50811) | 22(45218) |
| 20X | Normal | 58(97899) | 13(20610) | 12(32538) |
| 40X | Cancerous | 106(647960) | 23(172033) | 22(164026) |
| 40X | Normal | 58(374812) | 13(78663) | 12(126294) |

Supplementary Table 4: Individual performance for each subtype with Vanilla CNN in subtype classification

| Model | Subtype | Accuracy |
|---|---|---|
| Resnet-18 | KIRC | 97.09% |
| | KIRP | 69.75% |
| | KICH | 84.36% |
| Resnet-34 | KIRC | 97.35% |
| | KIRP | 72.49% |
| | KICH | 79.72% |

Supplementary Table 5: Individual performance for each subtype with CNN + DAGSVM in subtype classification

| Model | Subtype | Accuracy |
|---|---|---|
| Resnet-18 | KIRC | 94.84% |
| | KIRP | 77.25% |
| | KICH | 95.12% |
| Resnet-34 | KIRC | 91.30% |
| | KIRP | 82.41% |
| | KICH | 95.47% |

Supplementary Table 6: Classification of subtypes Normal tissues using model trained on Cancerous tissues

| Model | Precision | Recall | AUC |
|---|---|---|---|
| Resnet 18 | 79.88% | 79.85% | 0.85 |
| Resnet 34 | 79.44% | 79.12% | 0.85 |

Supplementary Table 7: KIRC Patient characteristics

| KIRC | Summary | Median |
|---|---|---|
| **Number of Patients** | 469 | |
| **Status** | | |
| Alive | 310 | |
| Dead | 159 | |
| **Age** | 26-90 years | 61.42 years |
| **Gender** | | |
| Male | 300 | |
| Female | 169 | |
| **Grade** | | |
| 1 | 10 | |
| 2 | 188 | |
| 3 | 175 | |
| 4 | 58 | |
| **Stage** | | |
| 1 | 228 | |
| 2 | 49 | |
| 3 | 117 | |
| 4 | 73 | |

Supplementary Table 8: Features extracted for Survival Analysis

| Tumor shape features | Cell Nuclei Features |
|---|---|
| Main region area | Total Area |
| Main region convex area | Total filled area |
| Main region filled area | Total convex area |
| Main region perimeter | Total Perimeter |
| Main region major axis | Total Minor axis |
| Total peri(squared) by area | Total Major Axis |
| Main region minor axis | Total peri(squared) by area |
| Main region peri(squared) by area | |
| Main region eccentricity | |
| Main region solidity | |
| Main region angle | |
| Total area | |
| Total convex area | |
| Total filled area | |
| Total major axis | |
| Total minor axis | |
| Total perimeter | |
| Total peri(squared) by area | |

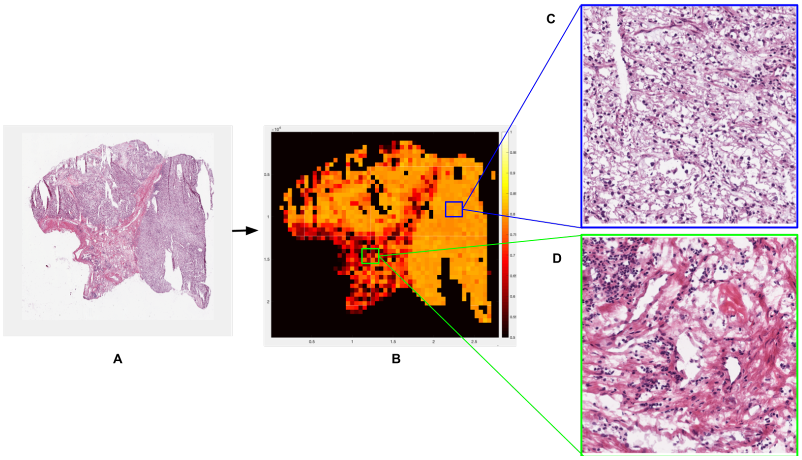


Figure 1: **Visualization of High probability regions of KIRC** - High probability regions (displayed in blue) and low probability regions (displayed in green)

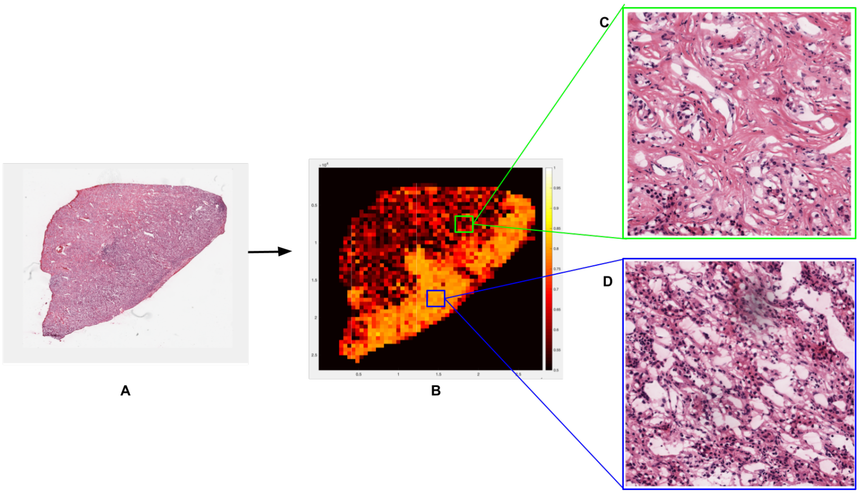


Figure 2: **Visualization of High probability regions of KIRC -** High probability regions (displayed in blue) and low probability regions (displayed in green)
